# The Na^+^/K^+^ ATPase Regulates Glycolysis and Modifies Immune Metabolism in Tumors

**DOI:** 10.1101/2020.03.31.018739

**Authors:** Sydney M. Sanderson, Zhengtao Xiao, Amy J. Wisdom, Shree Bose, Maria V. Liberti, Michael A. Reid, Emily Hocke, Simon G. Gregory, David G. Kirsch, Jason W. Locasale

**Affiliations:** Department of Pharmacology and Cancer Biology, Duke University School of Medicine, Durham, NC, 27710; Department of Radiation Oncology, Duke University School of Medicine, Durham, NC, 27710, USA; Duke Molecular Physiology Institute, Duke University School of Medicine, Durham, NC, 27710, USA

## Abstract

Cancer therapies targeting metabolism have been limited due to a lack of understanding of the controlling properties of vulnerable pathways. The Na^+^/K^+^ ATPase is responsible for a large portion of cellular energy demands but how these demands influence metabolism and create metabolic liabilities are not known. Using metabolomic approaches, we first show that digoxin, a cardiac glycoside widely used in humans, acts through disruption to central carbon metabolism via on target inhibition of the Na^+^/K^+^ ATPase that was fully recovered by expression of an allele resistant to digoxin. We further show in vivo that administration of digoxin inhibits glycolysis in both malignant and healthy cells, particularly within clinically relevant cardiac tissue, while exhibiting tumor-specific cytotoxic activity in an allografted soft tissue sarcoma. Single-cell expression analysis of over 31,000 cells within the sarcoma shows that acute Na^+^/K^+^ ATPase inhibition shifts the immune composition of the tumor microenvironment, leading to selective alterations to metabolic programs in specific immune cells thus acting both through tumor cell and microenvironmental (e.g. macrophage) cells. These results provide evidence that altering energy demands can be used to regulate glycolysis with cell-type specific consequences in a multicellular environment of biomedical interest.

## Introduction

Cancer cells exhibit metabolic reprogramming to support their uncontrolled proliferation, most notably in their utilization of processes that ultimately generate energy including glycolysis and the tricarboxylic acid (TCA) cycle (collectedly referred to as central carbon metabolism). Energy metabolism is coupled to redox, biosynthesis, and signaling^1,2^. Altered metabolism such as that observed in cancer has been attributed to a myriad of factors, such as oncogene activation^3–5^, dysregulation of mitochondria^6–9^, adaptation to oxygen^10,11^ and nutrient^12^ scarcity. While many of these alterations have been shown to promote dependencies on central carbon metabolism, therapies targeting these metabolic processes are toxic, and the efficacy of those that are sufficiently tolerable have poorly understood specificities^13,14^. Therefore, novel mechanisms that reveal vulnerabilities associated with metabolic reprogramming remain highly desired.

Cardiac glycosides (CGs) are tolerated agents commonly prescribed for the treatment of cardiac arrythmias or congestive heart failure. They are largely believed to act as inhibitors of the sodium-potassium pump (also referred to as the Na^+^/K^+^ ATPase)^15^. This transmembrane enzyme imports two potassium ions while exporting three sodium ions in an ATP-dependent manner, thereby maintaining the electrochemical gradient across the cell membrane^16^. The activity of this pump additionally contributes to the regulation of intracellular pH^17^, glucose uptake^18^, and Ca^2+^ levels^19^.

Interestingly, CGs have been shown to exhibit anticancer activity^20–22^. Nearly half a century ago, Efraim Racker postulated that a partially defective Na^+^/K^+^ ATPase could be a primary tumorigenic event by disrupting the cellular energetic state^23^. Given that the Na^+^/K^+^ ATPase accounts for roughly 20-70% of cellular ATP demand^24,25^, it was suggested that lowered enzyme activity could induce dysregulation of ATP-producing processes and resulting cellular programming towards an oncogenic state^23^. While this hypothesis has remained largely unexplored, it suggests that the Na^+^/K^+^ ATPase could be a major mechanism of control over central carbon metabolism.

In this study, we show that an immediate and direct consequence of Na^+^/K^+^ ATPase inhibition is disruption of central carbon metabolism and then show how this reprograms nearly all of metabolism. We demonstrate that the downstream metabolic consequences of digoxin treatment are specifically mediated through on-target inhibition of the Na^+^/K^+^ ATPase (that could be fully resotred by a resistant allele of Na^+^/K^+^ ATPase) and that digoxin treatment can impact metabolic processes in both healthy and malignant tissues with differential requirements in malignant cells. Furthermore, we use single-cell RNA sequencing to explore the metabolic consequences of Na^+^/K^+^ ATPase inhibition to show that acute inhibition reveals specific metabolic requirements in the tumor microenvironment and exerts intriguing metabolic changes in different immune cell compartments leading to coordinated regulation of non-metabolic genes.

## Results

### Digoxin disrupts central carbon metabolism and related processes in a time- and dose-dependent manner

To determine whether cell sensitivity to digoxin is associated with intrinsic metabolic state, we compared basal metabolic uptake and excretion rates with digoxin IC_50_ values across a panel of 59 cancer cell lines. This analysis demonstrated that digoxin treatment correlated with the metabolic flux of TCA intermediates (Figure 1A), including malate (Spearman correlation, *r* = −0.38, *p* = 0.0025) and citrate (Spearman correlation, *r* = −0.35, *p* = 0.0056), as well as the ATP-recycling metabolite creatine (Spearman correlation, *r* = 0.29, *p* = 0.02) (Figure S1A). This initial finding suggested that Na^+^/K^+^ ATPase activity could be linked to central carbon metabolic processes.

**Figure 1.**
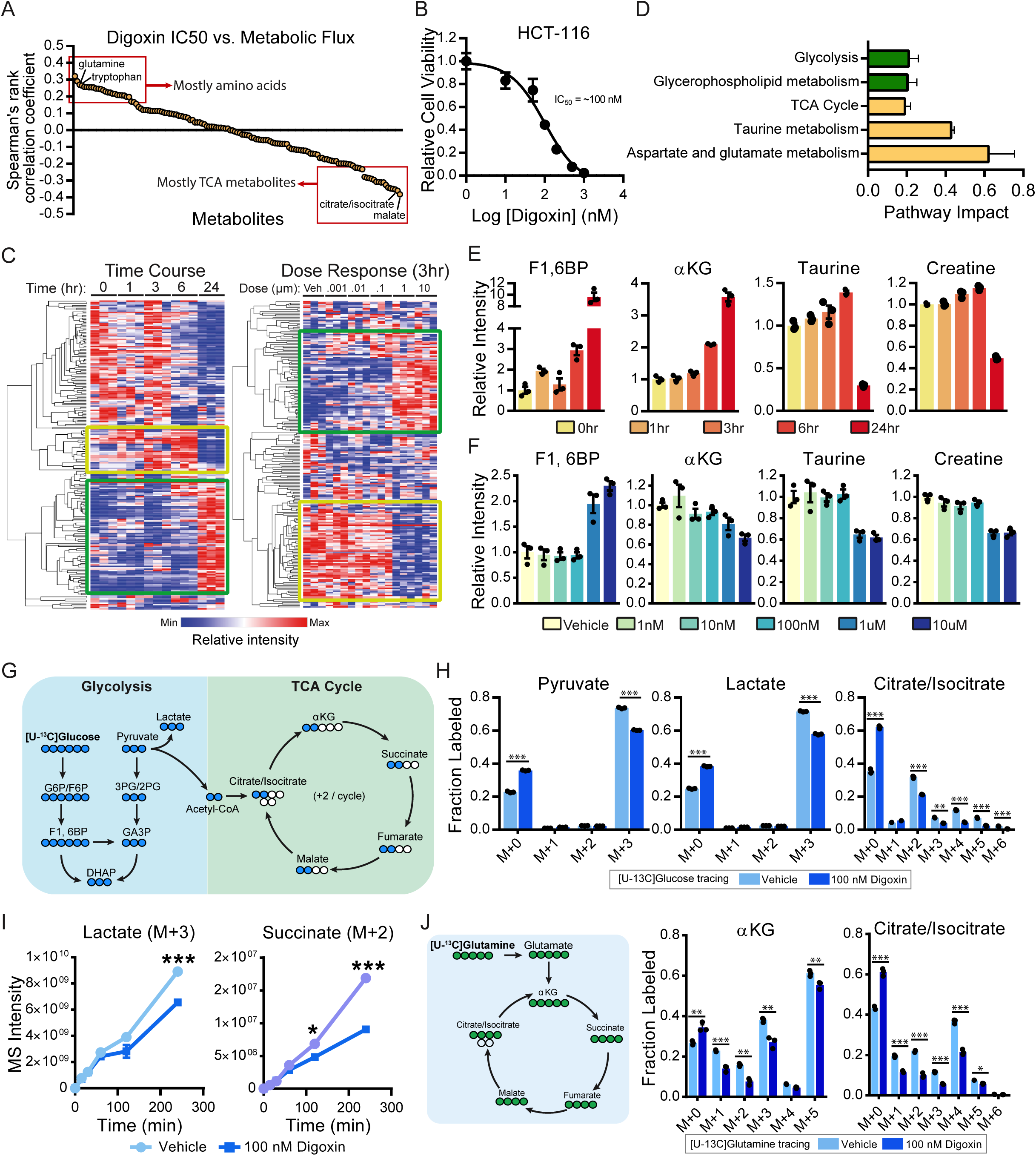
Digoxin disrupts central carbon metabolism and related processes in a time- and dose-dependent manner. (**A**) Spearman rank correlations of digoxin IC_50_ values and basal metabolic flux of individual metabolites in NCI-60 cancer cell panel. The metabolites are ranked based on the correlation coefficients. (**B**) Experimental measurement of digoxin IC_50_ in HCT-116 cells after 48 hours. (**C**) Clustered heatmaps of relative intensities of global metabolites upon digoxin treatment at indicated time points (left) and dose levels (right). Colored boxes highlight the most prominent metabolic changes. (**D**) Enriched metabolic pathways determined from significantly altered metabolites (p < 0.05, Student’s t-test) indicated in (C). The bar color matches box color in (C). (**E and F**) Relative intensities of representative central carbon and energy metabolites at different time points (E) and dose levels (F). (**G**) Diagram of isotopologue labeling of [U-^13^C] glucose through glycolysis into the TCA cycle. (**H**) Fractional abundance of each [U-^13^C] glucose labeled isotopologue relative to the sum of all isotopologues of the glycolytic and TCA intermediates. (**I**) Relative intensities of biologically relevant [U-^13^C] glucose labeled isotopologues of lactate and succinate at different time points. (**J**) Diagram of isotopologue labeling from [U-^13^C] glutamine to the TCA cycle (left) and fractional abundance of each [U-^13^C] glutamine-labeled isotopologue of TCA intermediates (right).

To assess the metabolic consequences of digoxin treatment, we generated metabolite profiles of HCT-116 cells both temporally using the digoxin IC_50_ concentration of 100 nM (Figure 1B), as well as acutely (3 hours) using increasing concentrations of digoxin (Figure S1B and 1C). Pathway analyses demonstrated that central carbon metabolism, as well as processes associated with its activity including aspartate/glutamate metabolism and taurine metabolism, were among the most significantly impacted metabolic pathways by digoxin in both a time- and dose-dependent manner (Figure 1D).

Closer examination of these metabolite profiles revealed an increase in upper glycolytic intermediates as well as a decrease in TCA cycle metabolites in both a time-(Figure 1E and S1C) and dose-dependent (Figure 1F and S1C) manner. Importantly, the accumulation of fructose 1,6-bisphosphate (F1,6BP) has been shown to indicate changes to glycolytic flux ^26^, indicating that the observed increases in F1,6BP levels (Figure 1E and 1F) are demonstrative of disruption to central carbon metabolism. Additionally, we consistently observed decreases in the levels of taurine (Figure 1F) and hypotaurine (Figure S1D), as well as creatine (Figure 1F) and phosphocreatine (Figure S1E). Taurine has been suggested to partially regulate mitochondrial electron transport chain (ETC) activity ^27^, while creatine enables the rapid anaerobic recycling of ATP under high energetic demand ^28^, thus providing additional evidence of disrupted energy status.

To study the effects of digoxin on downstream processes of central carbon metabolism, we first measured steady-state glucose incorporation into glycolysis and the downstream TCA cycle using media supplemented with uniformly labeled [^13^C]-glucose ([U-^13^C]-glucose) (Figure 1G and S1F). We found reductions in the labeling of both glycolytic and TCA intermediates (Figure 1H and S1G). Additional assessment of the kinetics of glycolytic flux into the mitochondria further revealed that the labeling of TCA intermediates [U-^13^C]-glucose administration was significantly reduced (Figure 1I and S1H). Finally, by measuring the steady-state incorporation of glutamine into the TCA cycle via conversion to α-ketoglutarate (α-KG) using [U-^13^C]-glutamine (Figure 1J), we found significantly reduced labeling of TCA intermediates (Figure 1J and S1I) further indicating alterations in mitochondrial activity. Altogether, these findings demonstrate that digoxin substantially disrupts multiple nodes of central carbon metabolism which extends to other intracellular energetic processes.

### Digoxin exerts its metabolic effects via on-target inhibition of the Na^+^/K^+^ ATPase

Upon consideration of the metabolic reprogramming induced by digoxin treatment, we assessed whether the cytotoxic effects of digoxin could be mitigated by nutrient supply. We cultured HCT-116 cells in media containing 100 nM digoxin as well as supplementations of nutrients related to central carbon metabolism or redox balance which is coupled to central carbon metabolism; surprisingly, we found that the majority of these treatments were insufficient to rescue cells from digoxin treatment (Figure 2A), although the supplementation of either nucleosides or the antioxidant NAC were modestly cytoprotective (Figure 2B), in line with previous reports ^29^. These results suggest that the metabolic disruption induced by digoxin likely exerts extensive downstream consequences on cellular homeostasis and function beyond the targeting of a single pathway.

**Figure 2.**
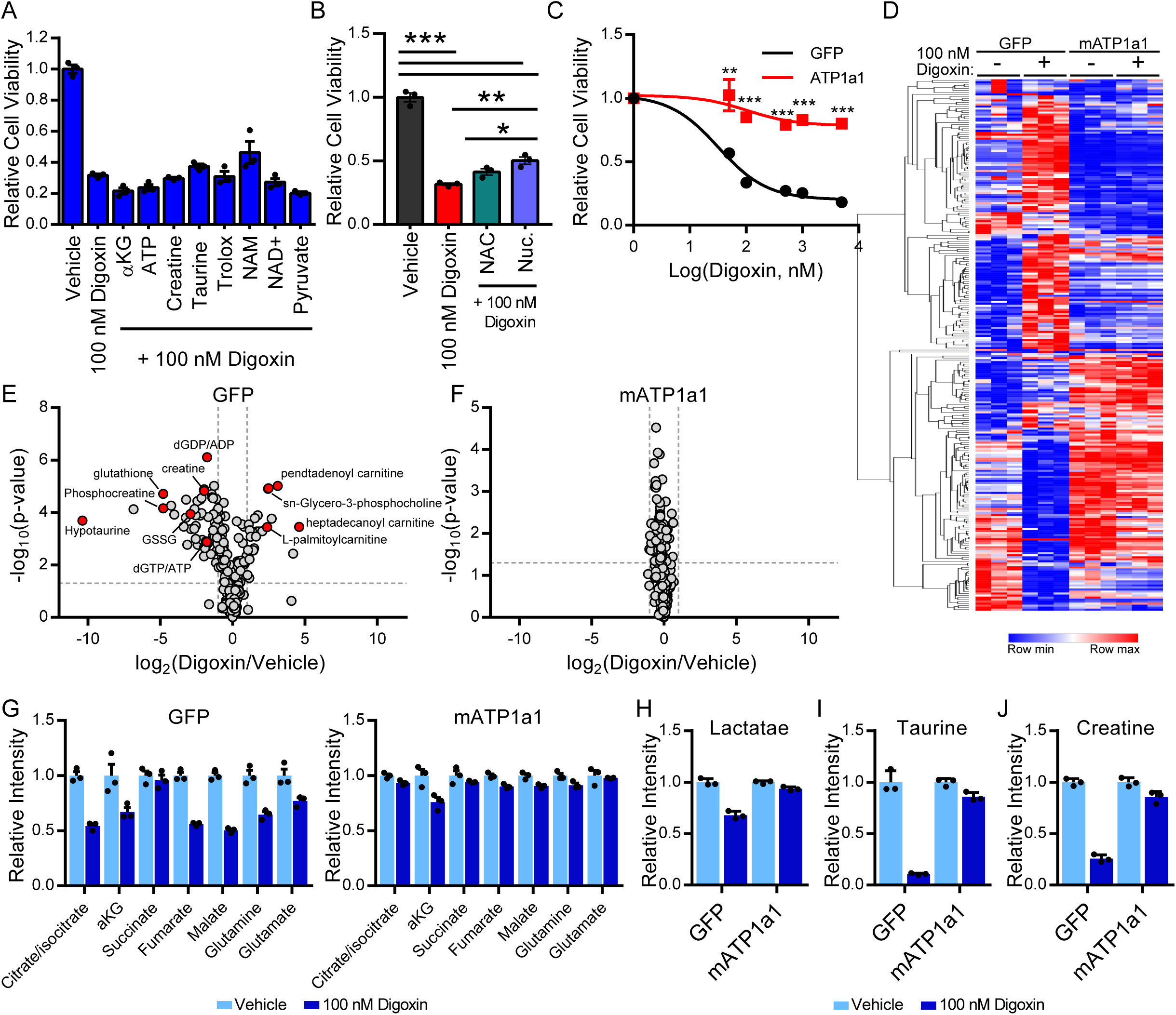
Digoxin exerts its metabolic effects via on-target inhibition of the Na^+^/K^+^ ATPase. (**A and B**) Relative cell viabilities of HCT-116 cells treated with vehicle or 100 nM digoxin in regulator media or with supplementation of the indicated nutrient for 72 hours. Nuc.: nucleosides. (**C**) Relative cell viability of HCT-116 cells transfected with mATP1a1 digoxin-resistant subunit, cultured in incremental concentrations of digoxin. (**D**) Heatmap of fold changes in global metabolite levels with or without 48-hours 100 nM digoxin treatment in HCT-116 transfected cells. (**E and F**) Volcano plots displaying metabolites profiles of HCT-116 cells transfected with GFP control (E) or mATP1a1 (F) vectors after treated with digoxin and vehicle. (**G—J**) Relative intensities of TCA cycle metabolites (G), lactate (H), taurine (I) and creatine (J) of HCT-116 cells transfected with GFP or mATP1a1 vectors upon digoxin and vehicle treatment.

The possibility remained that the metabolic consequences of digoxin were due to factors other than inhibition of Na^+^/K^+^ ATPase activity. It is established that murine Na^+^/K^+^ ATPase enzymes are substantially less sensitive to CGs, with the mouse isoform exhibiting roughly 1000-fold lower affinity for CGs than the human counterpart ^30,31^. Indeed, it has been shown that ectopic expression of the α-subunit of the mouse ATPase (mATP1a1) is sufficient to rescue cell viability upon digitoxin treatment ^32^. Therefore, to determine whether the observed metabolic effects were specifically attributable to disruption of Na^+^/K^+^ ATPase activity, we ectopically expressed the mouse mATP1a1 subunit in HCT-116 cells and demonstrated a complete rescue of cell viability after treatment with digoxin (Figure 2C). We then treated the mATP1a1-expressing cells with digoxin at the IC_50_ concentration and compared the resulting metabolite profiles (Figure 2D). We found that mATP1a1 expression completely blocked the metabolic consequences of digoxin treatment (Figure 2E and 2F), thereby restoring the activity of central carbon metabolism (Figure 2G and 2H) as well as taurine (Figure 2I) and creatine (Figure 2J) levels. This finding establishes that disrupted central carbon metabolism is an intrinsic component of digoxin-induced cytotoxicity, which is a direct result of Na^+^/K^+^ ATPase inhibition.

### Digoxin treatment impacts energy metabolism in a tissue-specific and antineoplastic manner

After defining the impact of digoxin on central carbon metabolism and related energetic processes in cell culture, we next sought to determine whether these effects could be achieved in a more complex *in vivo* setting. Numerous findings of CGs effectively inhibiting tumor growth in xenograft studies have been reported ^29,32,33^; however, given the extreme (greater than 1000-fold) differential in CG-binding to the ATPase enzymes found in human-derived cells and the murine host, these findings are confounded by the artificially high concentrations in xenograft studies of digoxin that can be used with no effects on the mouse host due to their weak binding affinity to murine Na^+^/K^+^ ATPases. Thus, a therapeutic window needed to achieve tumor growth inhibition could be prohibitive in a setting where the host and tumor express the same ATPase enzymes ^34^. Additionally, it remains to be explored whether Na^+^/K^+^ ATPase inhibition can impact metabolism in both healthy and malignant tissue.

To investigate these essential considerations, we cultured mouse sarcoma cells generated from primary sarcoma tumors driven by expression of oncogenic KRAS^G12D^ and p53 deletion (*Kras*^*LSL-G12D/+*^;*Trp53*^*flox/flox*^, or KP) and found that they exhibited an IC_50_ of 100 μM, in line with their murine isoform expression (Figure S2A). As proof of principle, we first verified that administration of this concentration reliably impacted central carbon metabolism in cell culture (Figure S2B and S2C). We then orthotopically injected these cells into the right gastrocnemius muscle of syngeneic 129/SvJae mice. Upon tumor palpation (approximately 11 days after injection), we treated mice with a previously reported dose^35^ of 2 mg/kg digoxin every 24 hours and collected tumors as well as healthy tissues after administration of the fourth dose (Figure 3A). Of note, while this treatment regimen appeared to trend towards tumor growth inhibition, treated mice exhibited some weight loss upon daily digoxin treatment (Figure S2D and S2E) although no other physiological or behavioral signs of toxicity were observed.

**Figure 3.**
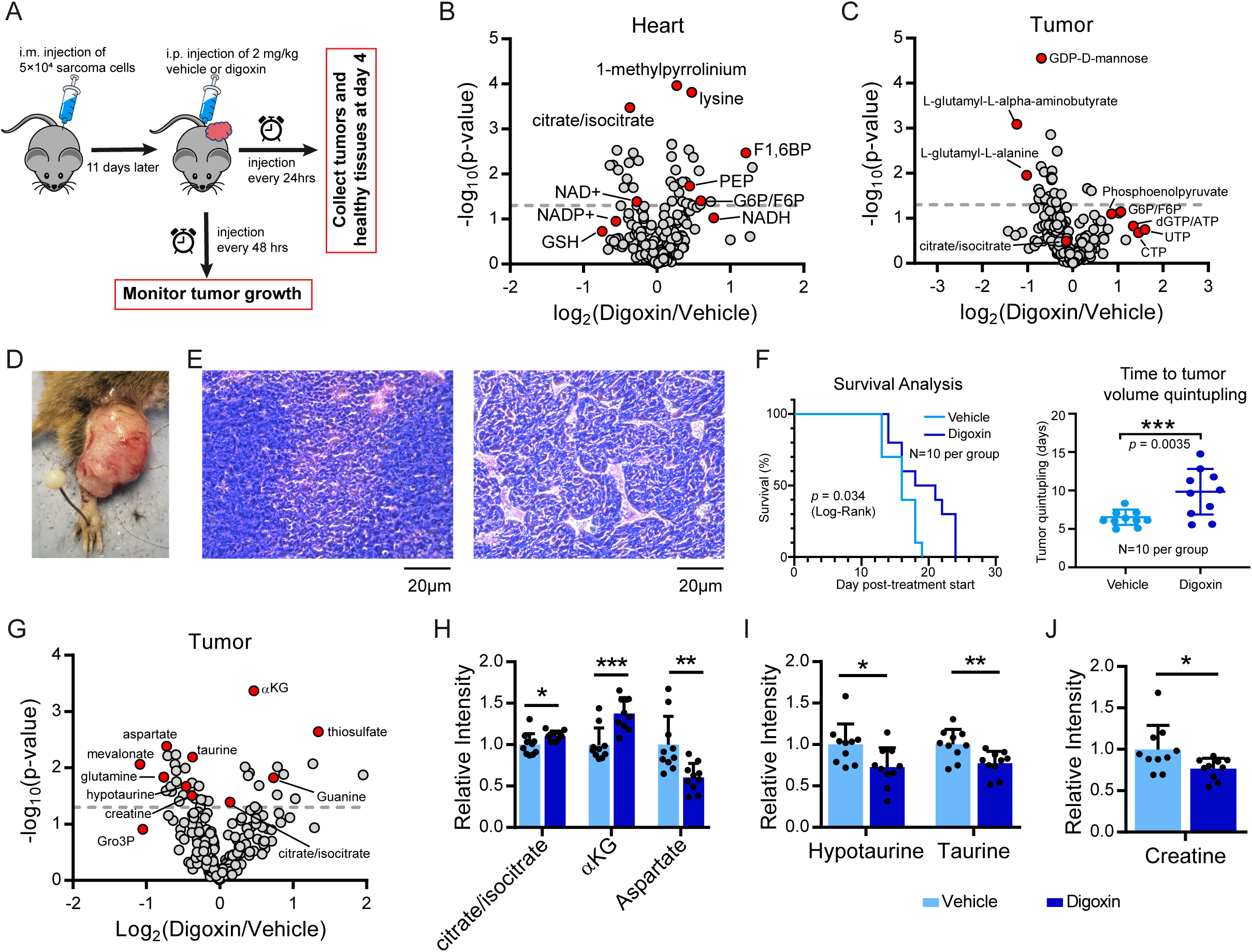
Digoxin treatment impacts energy metabolism in a tissue-specific and antineoplastic manner. (**A**) Diagram of digoxin treatment schedule in allograft sarcoma model. i.m., intramuscular; i.p., intraperitoneal injection. (**B**) Volcano plots displaying metabolites profiles of cardiac tissue between vehicle and digoxin treatment. (**C**) Same as in (B) but for tumor tissue. (**D**) Representative image of allograft sarcoma at growth endpoint. The tumor measured 12.8 mm ×13.4 mm at the time it was determined to have reached the endpoint. (**E**) H&E sections of representative tumors in vehicle treatment, showing regions of both low (left) and high (right) vascularity. (**F**) Kaplan-Meier survival curve (left) and quantification of time for tumors to quintuple in volume (right). Values in the right panel are represented as mean ± SD. (**G**) Same as in (B) but for tumors exposed to chronic digoxin treatment. (**H—J**) Relative intensities of TCA intermediates (H), taurine metabolites (I), and creatine (J) in chronic vehicle- and digoxin-treated tumors. N=10.

Metabolite profiling revealed that cardiac tissue was the most significantly impacted tissue type (Figure 3B), with substantial alterations in glycolytic (Figure S2F) and TCA cycle (Figure S2G) intermediates. These effects were even more pronounced in cardiac than tumor tissue in this context (Figure 3B and 3C). A number of similar changes were also found in muscle, brain, liver, and kidney tissue (Figure S2H), although to a much lesser extent than was found in cardiac tissue (Figure S2F, S2G, S2I and S2J) or the cultured sarcoma cells (Figure S2B and S2C).

To examine the antineoplastic potential of digoxin in this setting, we treated the second cohort of orthotopically-engrafted syngeneic mice with digoxin (2mg/kg) or vehicle every 48 hours and monitored tumor growth (Figure 3D). There were no discernible histological differences between vehicle- and digoxin-treated groups, with the majority of tumors exhibiting regions of both low (Figure 3E, left) and high (Figure 3E, right) vascularity. Of note, this heterogeneous landscape, likely resulting from the relatively large size of these KP allograft tumors (Figure 3D), closely resembles the regional heterogeneity found in patient sarcomas^36^. Regression analyses demonstrated that digoxin treatment significantly delayed tumor growth as measured by time to tumor quintupling (Figure 3F). Furthermore, the most prominent metabolic alterations found in tumors exposed to chronic digoxin treatment were consistent with dysregulation of energy-related metabolic processes (Figure 3G), with disruptions to mitochondrial metabolism (Figure 3H) as well as taurine (Figure 3I) and creatine (Figure 3J) levels. Collectively, these findings demonstrate that digoxin can impact metabolic processes in both healthy and malignant tissues and that these metabolic perturbations lead to antineoplastic effects in a physiological setting.

### Acute digoxin treatment remodels the immune compartment of the tumor microenvironment

An important feature of allograft models is the presence of an intact immune system^37,38^. Given that the differential requirements of central carbon metabolism have become increasingly appreciated in immune cell function^39–41^, we performed single-cell RNA sequencing (scRNA-seq) to explore the effects of digoxin treatment on the tumor microenvironment. Briefly, we treated mice with vehicle or 2 mg/kg digoxin (using two mice per group) with the daily treatment regimen described previously (Figure 4A) and harvested the tumors after the final fourth dose. We then dissociated the tumors into single-cell suspensions and performed 10x scRNA-seq without cell type sorting or purification using the Chromium drop-seq platform (10x Genomics) (Figure 4A). We analyzed mRNA expression from >7,200 cells from each tumor after quality controls (Figure 4A), with roughly 100,000 reads from each cell that covered nearly 5,000 genes (Figure S3A—S3C), and used previously published scRNA-seq data of healthy murine muscle tissue^42^ as the reference to distinguish between malignant and non-malignant cells based on relative gene copy number^43^ (Figure 4B, S3D and S3E). Acute digoxin treatment trended towards a reduction in the relative population of malignant cells (Figure 4C), consistent with previous observations of early tumor growth inhibition (Figure S2D).

**Figure 4.**
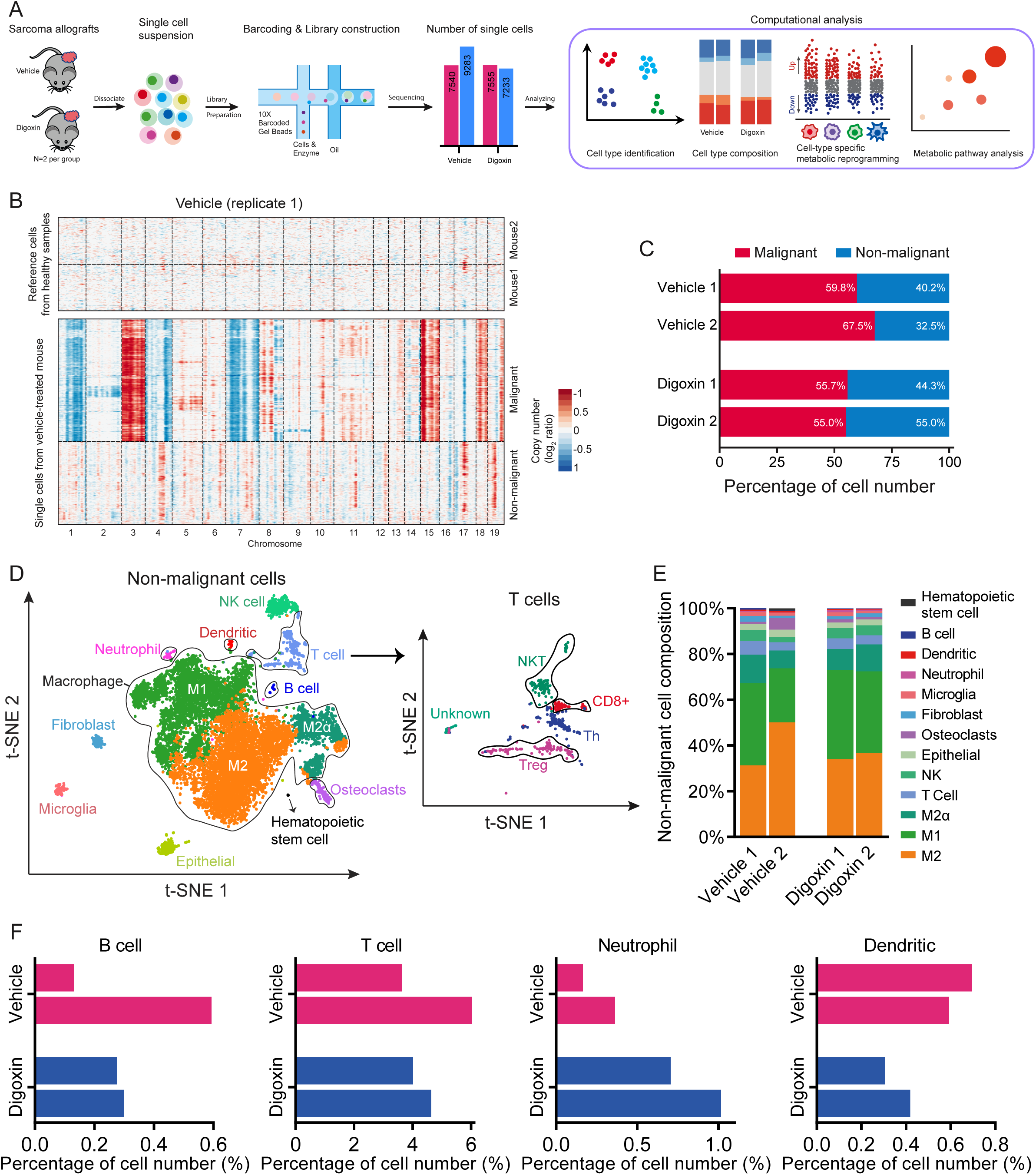
Acute digoxin treatment shifts the tumor microenvironment. (**A**) Overview of single-cell RNA sequencing workflow, from tumor generation, cell harvest, RNA sequencing, to gene expression analysis. (**B**) Chromosomal landscape of large-scale CNVs for individual cells (rows) from normal mouse muscle tissue and representative tumor vehicle replicate 1, allowing us to distinguish cells into malignant and non-malignant. Amplifications (red) and deletions (blues) were inferred by averaging gene expression over 100 genes stretch on each chromosome (columns). (**C**) Relative proportion of malignant and non-malignant cells in each treatment. (**D**) t-SNE plots show identified non-malignant cell populations (left) and T cell subpopulations (right). M1: type I macrophages; M2: type II macrophages; M2α: highly proliferation M2; NK: Natural killer cells; Tregs: Regulatory T cells; Th: T-helper cells; NKT: Natural killer T cells. (**E**) Relative proportion of cell populations in the total non-malignant cell pool. (**F**) Comparison of relative proportions of indicated cell populations in vehicle and digoxin treatment.

Upon further analysis of the single-cell transcriptomes, we identified fifteen immune cell populations, including five distinct T-cell populations (Figure 4D, S4A and S4B). Myeloid cells (i.e. type I and type II macrophages, M1 and M2 respectively) were the most abundant immune cell population (Figure 4E), consistent with previous reports of autochthonous KP sarcomas ^44^. While the relative populations of intratumoral B cells and T cells did not appear to be appreciably skewed by this short-term digoxin treatment, the relative populations of neutrophil and dendritic cell infiltrates were slightly increased and decreased, respectively (Figure 4F). These results demonstrate that acute digoxin treatment induces an immediate shift in the tumor microenvironment.

### Targeting the Na^+^/K^+^ ATPase with digoxin alters metabolic processes in tumor cells and immune infiltrates

To determine whether digoxin treatment exerts differential effects on metabolic programming210 between cell types, we compared the scRNA-seq transcriptome of each tumor cell population after digoxin treatment with vehicle treatment. Oxidative phosphorylation was the most extensively altered metabolic program in malignant cells, with significant increases in expression of Atp5k transcripts corresponding to the ATP synthase gene (Figure 5 A and B). ATP synthase expression is linked to glycolytic activity, thereby providing additional evidence that central carbon metabolism disruption is an immediate and direct consequence of digoxin treatment. Further functional enrichment analysis of all significantly altered genes showed that the apoptosis process was activated (Figure S5), which could be induced by mitochondrial OXPHOS system^45,46^.

**Figure 5.**
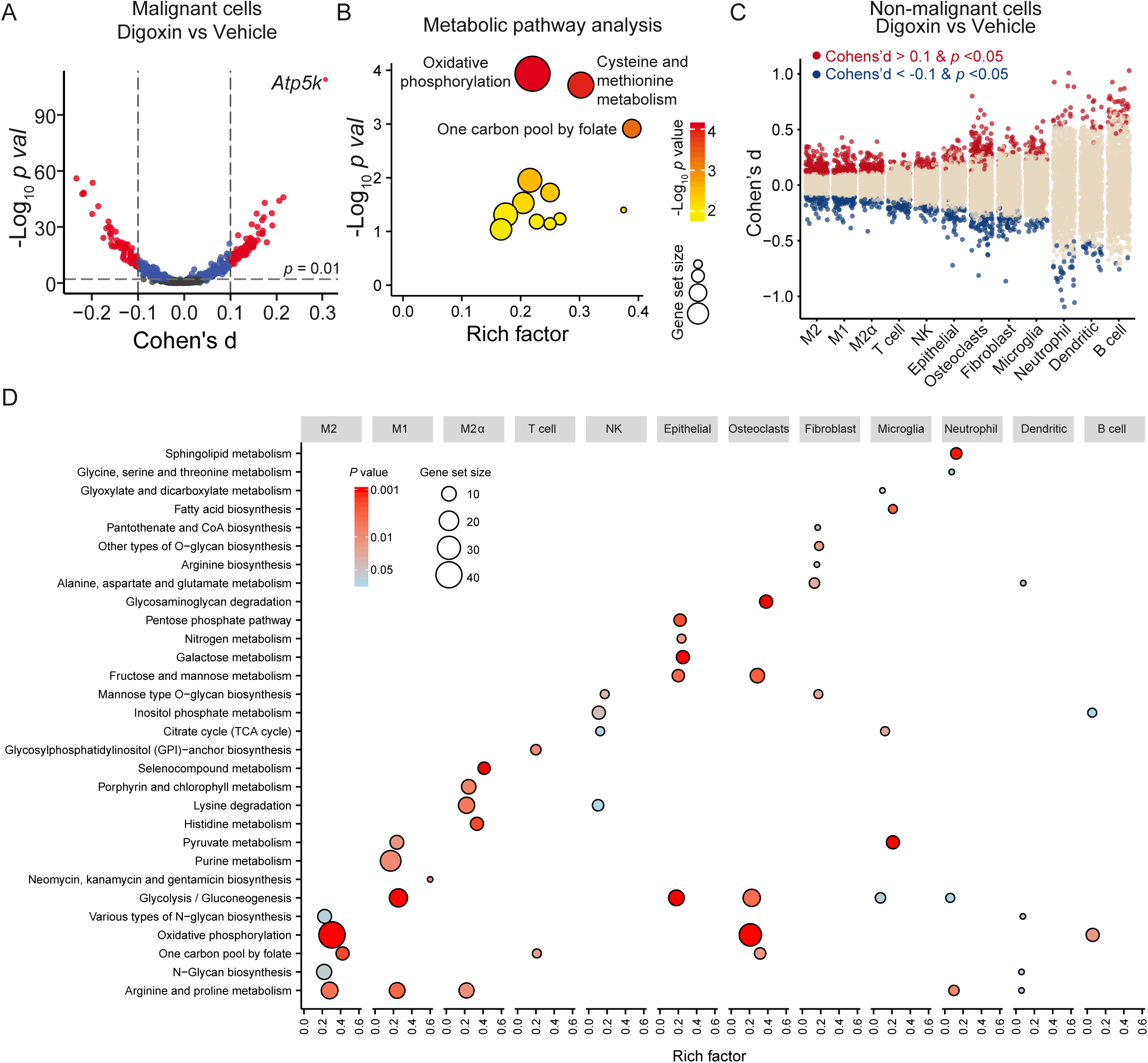
Digoxin treatment is associated with transcriptional reprogramming of metabolic processes in malignant and non-malignant cells. (**A**) Volcano plot of significance (measured by *p* value) against magnitude (measured by Cohen’s d) of metabolic gene expression differences between vehicle and digoxin treatment. The differential genes above a significance threshold of *p* value < 0.01 and the absolute value of Cohen’s d > 0.1 are labeled in red. (**B**) Metabolic pathways enriched in pathway analysis using differentially expressed metabolic genes determined from (A). Rich factor is the ratio of the number of enriched genes (represented by the size of the dots) to the number of background genes in the corresponding pathway. (**C**) Distribution of Cohen’s d for metabolic genes in each non-malignant cell population, with significantly altered genes (p < 0.05 and | Cohens’d | > 0.1) marked in red (upregulated in digoxin treatment) or blue (downregulated in digoxin treatment). (**D**) Enriched metabolic pathways in non-malignant cell populations, determined by differentially expressed metabolic genes in (C). For each cell population, the top 3 enriched pathways are shown.

Upon examination of the non-malignant cell populations, we found that different cell populations were subject to various magnitude alterations on their metabolic genes after digoxin treatment (Figure 5C). Pathway analysis of the significantly altered metabolic genes (|Cohens’ d| > 0.1 and p-value < 0.05) suggested that digoxin exposes diversity in metabolic plasticity among different cell populations (Figure 5D and S6). The macrophage populations exhibited metabolic reprogramming specifically within central carbon metabolism, with M1 macrophages characterized by upregulation of glycolysis while M2 macrophages were characterized by an upregulation in oxidative phosphorylation similar to malignant cells (Figure 5D). Interestingly, previous studies of macrophage behavior have also observed similar metabolic shifts in these two populations upon their activation^47,48^. To examine this possibility more closely, we performed KEGG pathway analysis on their full transcriptomes and additionally found transcriptional signatures (i.e. endoplasmic reticulum associated genes) consistent with their polarization^49–51^ (Figure S7 A-C). These observations illustrate that digoxin is associated with broad transcriptional consequences, including within metabolic processes, across multiple tumor cell populations.

## Discussion

Metabolic programming, on both a cellular and physiological level, is known to be highly regulated by environmental factors which are extrinsic to discrete metabolic reactions^52–56^. In line with this, many conventional therapies have been shown to exert substantial metabolic effects beyond their understood mechanism of action^57–59^. As many of the metabolic vulnerabilities inherent to cancer cells are difficult to target without inducing toxic consequences on healthy tissue, the identification of agents that can be repurposed to exploit these processes remains an active area of investigation^60–63^.

Cardiac glycosides have been shown to exhibit antineoplastic effects in numerous settings^20,21,35^, which have been attributed to a myriad of sources such disruption of proton gradients^32^ and activation of kinases that physically interact with the sodium-potassium pump (i.e. Na^+^/K^+^ ATPase “signalosome”)^64–66^. However, fluctuations in energetic demand from membrane transport activity have been shown to impact glycolytic rate^67^, and it has been historically hypothesized that alterations in Na^+^/K^+^ ATPase activity may be a contributing factor to enhanced dependence on glycolysis in cancer^23^. Our results demonstrate that digoxin, through on-target inhibition of the Na^+^/K^+^ ATPase, induces broad metabolic disruptions in a time- and dose-dependent manner; these disruptions were most prominent within central carbon and energy-related metabolic processes, including taurine and creatine metabolism. Interestingly, taurine has been suggested to act as a mild cardiac glycoside in cardiac tissue through its modulation of mitochondrial ROS^27^, and taurine loss has been associated with Na^+^ efflux thereby reducing the degree of Ca^2+^ overload^68^. It’s therefore tempting to speculate that the dual loss of taurine and creatine upon digoxin treatment are compensatory responses to ion imbalance and reduced ATP production, respectively.

Our results further provide the first characterization of the global metabolic consequences of digoxin on healthy tissues to allow direct comparison with metabolic profiles in tumors. We found that acute digoxin treatment effectively impacted central carbon metabolism or other energy-related metabolites (i.e. taurine and creatine) in diverse healthy tissues, with the most observable metabolic consequences found in cardiac tissue. Although the higher degree of central carbon metabolic disturbance in cardiac compared to tumor tissue after short-term digoxin treatment was unexpected, the use of digoxin for treating cardiac dysfunction likely indicates a tissue-specific affinity for CGs; indeed, it has been shown that the human cardiac-specific alpha subunit isoform exhibits a higher affinity for CGs compared to the isoforms present in most other tissues^69^, thereby providing a potential mechanism for the observed effects. Importantly, despite these prominent metabolic alterations in cardiac tissue, we did not observe any signs of cardiotoxicity at the doses we used, and these alterations were considerably more prominent in tumors with the antineoplastic long-term digoxin treatment, indicating the enhanced dependency of tumor cells on these processes.

We have recently demonstrated that scRNA metabolic gene transcriptomes can be used to investigate aspects of metabolic activity and plasticity within individual cells ^46^. Our results additionally contribute to our understanding of unique cell-autonomous responses within the tumor microenvironment by using scRNA-seq to examine diverse cell populations and metabolic programs within tumors following acute cytotoxic treatment. Our findings of both altered representations of immune infiltrate, as well as transcriptional programs of metabolic processes featuring clear distinctions between malignant and immune cell populations, in response to short-term digoxin treatment provide a novel glimpse into cell-specific intratumoral heterogeneity.

It remains unclear whether the altered metabolic programs and possible activation of different myeloid populations are a direct result of exposure to digoxin or indirect response to signals from adjacent malignant cells; furthermore, the observed variations in cell populations might become more pronounced with longer exposure to digoxin or at different stages of tumor development. Indeed, the potential dual polarization of M1 and M2 macrophages (which are commonly characterized as pro-inflammatory or anti-inflammatory, respectively)^70^ is intriguing and warrants further investigation. It will be interesting in future studies to determine the functional consequences of these variations in metabolic reprogramming between immune cell populations, especially upon consideration of previous studies reporting consequences of digoxin on immune cell activity in various noncancerous settings ^71,72^. Additional studies of these interactions could illuminate possible synergistic mechanisms that may elicit enhanced clinical efficacy of digoxin when combined with targeted immune therapies as has been found with other compounds^44,73,74^.

## Methods

### Cell Culture

Cells were cultured at 37°C, with 5% atmospheric CO2 in RPMI-1640 (Gibco), 10% heat-inactivated fetal bovine serum (FBS; F2442, Sigma), 100 U/mL penicillin (Gibco), and 100 mg/mL streptomycin (Gibco). HCT-116 cells were obtained from the American Tissue Culture Collection (ATCC), and cultured murine sarcoma cells were generated from primary murine sarcomas described previously ^44^ and below in “Allograft Mouse Studies”.

### Digoxin IC_50_ measurements

Cells were plated at a density of 5.0×10^3^ cells/well in triplicate in a 96-well plate and were allowed to adhere in full RPMI 1640 media for 24 hours. Cells were briefly washed with PBS and then incubated in medium containing vehicle (DMSO; #97061-250, VWR) or indicated concentrations of digoxin (#D6003, Sigma Aldrich). After 48 hours, the media was aspirated and replaced with 100 μl phenol-red free RPMI-1640 (Gibco) and 12 mM 3-[4,5-Dimethylthiazol-2-yl]-2,5-diphenyltetrazolium (MTT) (Thermo Fisher Scientific, #M6494). After 4 hours, the MTT-containing media was aspirated and 50 μl DMSO was added to dissolve the formazan. After 5 minutes, absorbance was read at 540 nm. For all experiments, three technical replicates per culture condition were used.

### Nutrient Rescue Experiments

For all nutrient rescue experiments, cells were plated at a density of 5.0×10^5^ cells/well in triplicate in a 6-well plate and were allowed to adhere for 24 hours. Cells were briefly washed with PBS and then incubated with vehicle (DMSO) or 100 nM digoxin, as well as one or more of the following supplementations: 5mM N-acetyl-cysteine (NAC; A9165, Sigma), 2 μM Trolox (238813, Sigma), 100 μM adenosine 5’-triphosphate disodium salt hydrate (ATP; A26209, Sigma), 10mM creatine (C3630, Sigma), 10mM Taurine (T8691, Sigma), 2μM dimethyl-a-ketoglutarate (cell-permeable a-KG; 349631, Sigma), 5mM sodium pyruvate (sc-208397A, Santa Cruz Biotechnology), 500 μM nicotinamide (N0636, Sigma), 100μM beta-nicotinamide adenine dinucleotide (NAD^+^; 160047, MP Biomedical), and 1 × Embryomax Nucleosides (ES-008-D, Millipore). For glucose-restricted medium, RPMI 1640 glucose- and glutamine-free medium was supplemented with 2 mM glutamine, 10% heat-inactivated FBS, 100 U/mL penicillin, and 100 mg/mL streptomycin; indicated concentrations of glucose were then serially added to the medium. After 48 hours, an MTT cell viability assay was performed as described above. For all experiments, three technical replicates per culture condition were used.

### [U-^13^C] glucose and [U-^13^C] glutamine Tracing

[U-^13^C] glucose (CLM-1396-10) and [U-^13^C] glutamine (CLM-1822-H-PK) were purchased from Cambridge Isotope Laboratories. [U-^13^C] glucose was added to RPMI 1640 glucose-free medium at a concentration of 11 mM, while [U-^13^C] glutamine was added to glucose- and glutamine-free RPMI-1640 medium (supplemented with 11 mM glucose) to a concentration of 2 mM. Cells were plated at a density of 3.0 × 10^5^ cells/well in a 6-well plate, and after 24 hours were treated with either vehicle (DMSO) or 100 nM digoxin. After 4 hours of drug treatment, the medium was quickly aspirated and washed with glucose-free RPMI 1640 medium, and either [U-^13^C] glucose- or [U-^13^C] glutamine-containing medium was added to each well. Cells were then collected for metabolite extraction either 20 hours later or at the indicated time points. For all experiments, three technical replicates per culture condition were used.

### Lentiviral Transfection and Transduction

HEK-293T cells were plated at a density of 1.0 × 10^6^ cells/10 cm plate in RPMI 1640 (Gibco) supplemented with 10% heat-inactivated FBS, penicillin (100 U/ml), and streptomycin (100 mg/ml) and were allowed to adhere and reach 70% confluency. 15 μg of mATP1a1 (EX-Mm01329-Lv105, GeneCopoeia) or GFP control (EX-EGFP-Lv105, GeneCopoeia) plasmid, 10 μg of PsPAX2 packaging vector (no. 12260, Addgene), and 5 μg of PMD2.G envelope– expressing plasmids (no. 12259, Addgene) were diluted in 500 μl of jetPRIME buffer (no. 114-07, Polyplus-transfection) and vortexed. Next, 60 μl of the jetPRIME transfection reagent (no. 114-07, Polyplus-transfection) was added to the mixture, vortexed for 10 seconds, and left to incubate for 10 minutes at room temperature. The medium in the plate was replaced with fresh medium, and the transfection mix was then added to the 10-cm plate dropwise. After 24 hours, the transfection medium was replaced with fresh medium. After an additional 24 hours, the medium was collected and filtered through a 0.45-μm filter for virus collection. HCT-116 cells were plated in 10-cm plates, and when they reached 30-50% confluency, virus-containing medium (1:1 with fresh RPMI 1640 medium) was added to the plates along with polybrene (4 μg/μl). After 24 hours, the virus-containing medium was removed and replaced with fresh RPMI 1640 medium. Cells were incubated with puromycin (1 μg/ml) for 48 hours for positive selection.

### Metabolite Extraction

Media was quickly aspirated and 1 mL of extraction solvent (80% methanol/water, cooled to - 80°C) was added to each well of the 6-well plates, and were then transferred to −80°C for 15 minutes. Plates were removed and cells scraped into the extraction solvent on dry ice. All metabolite extracts were centrifuged at 20,000g at 4°C for 10 min. Finally, the solvent in each sample was evaporated in a speed vacuum, and the resulting pellets were stored in −80°C until resuspension. For polar metabolite analysis, the cell extract was dissolved in 15 μL water and 15 μL methanol/acetonitrile (1:1, v/v) (LC-MS optima grade, Thermo Fisher Scientific). Samples were centrifuged at 20,000g for 2 minutes at 4°C, and the supernatants were transferred to liquid chromatography (LC) vials.

### Liquid Chromatography

Ultimate 3000 HPLC (Dionex) with an Xbridge amide column (100 x 2.1 mm i.d., 3.5 μm; Waters) was coupled to Q Exactive-Mass spectrometer (QE-MS, Thermo Scientific) for metabolite separation and detection at room temperature. The mobile phase A reagent was composed of 20 mM ammonium acetate and 15 mM ammonium hydroxide in 3% acetonitrile in HPLC-grade water (pH 9.0), while the mobile phase B reagent was acetonitrile. All solvents were LC-MS grade and were purchased from Fischer Scientific. The flow rate used was 0.15 mL/min from 0-10 minutes and 15-20 minutes, and 0.3 mL/min from 10.5-14.5 minutes. The linear gradient was as follows: 0 minutes 85% B, 1.5 minutes 85% B, 5.5 minutes 35% B, 10 minutes 35% B, 10.5 minutes 25% B, 14.5 minutes 35% B, 15 minutes 85% B, and 20 minutes 85% B.

### Mass Spectrometry

The QE-MS is outfitted with a heated electrospray ionization probe (HESI) with the following parameters: evaporation temperature, 120°C; sheath gas, 30; auxiliary gas, 10; sweep gas, 3; spray voltage, 3.6 kV for positive mode and 2.5 kV for negative mode. Capillary temperature was set at 320°C and S-lens was 55. A full scan range was set at 60 to 900 (m/z), with the resolution set to 70,000. The maximum injection time (max IT) was 200 ms. Automated gain control (AGC) was targeted at 3,000,000 ions.

### Peak Extraction and Metabolomics Data Analysis

Data collected from LC-Q Exactive MS was processed using commercially available software Sieve 2.0 (Thermo Scientific). For targeted metabolite analysis, the method “peak alignment and frame extraction” was applied. An input file (“frame seed”) of theoretical m/z (width set at 10 ppm) and retention time of ~260 known metabolites was used for positive mode analysis, while a separate frame seed file of ~200 metabolites was used for negative mode analysis. To calculate the fold changes between different experimental groups, integrated peak intensities generated from the raw data were used. Hierarchical clustering and heatmaps were generated using Morpheus software (The Broad Institute, https://software.broadinstitute.org/morpheus/). For hierarchical clustering, Spearman correlation parameters were implemented for row and column parameters. Pathway enrichment analysis was conducted by MetaboAnalyst 3.0 software (http://www.metaboanalyst.ca/faces/home.xhtml) using HMDB IDs of the metabolites that were significantly enriched (p < 0.05). The pathway library used was *Homo sapiens* and Fishers’ Exact test was employed for over-representation analysis. Other quantitation and statistics were calculated using Graphpad Prism software.

### Allograft Mouse Studies

All animal studies were performed following protocols approved by the Duke University Institutional Animal Care and Use Committee (IACUC) and adhere to the NIH Guide for the Care and Use of Laboratory Animals. KP sarcoma cells were obtained from a primary *Kras*^*LSL-G12D/+*^;*Trp53*^*flox/flox*^ sarcoma. The tumor was dissected from the hind limb and dissociated by shaking for 45 minutes at 37°C in collagenase Type IV (Gibco), dispase (Gibco), and trypsin (Gibco). Cell suspension was then strained through a 40 µm filter, washed in PBS, and plated for culture. KP cells were maintained *in vitro* in DMEM (Gibco) containing 10% heat-inactivated FBS (Gibco), and 1% Penicillin/Streptomycin (Gibco) for 8-10 passages before transplanting into syngeneic mice. All mice were maintained on a pure 129/SvJae genetic background. For allograft tumor initiation, cultured KP murine cells were suspended in DMEM medium at a concentration of 5×10^6^ cells/mL, and 5×10^4^ cells were injected into the gastrocnemius muscle of recipient mice. When tumors reached 70-150 mm^3^ (as determined by caliper measurement in two dimensions), the sarcomas were randomized to vehicle or digoxin groups. Mice were administered an intraperitoneal (i.p.) injection of either vehicle (PBS) or 2 mg/kg digoxin (prepared in DMSO and then diluted in PBS) with a volume not exceeding 250 μL. For short-term treatments, injections were administered every 24 hours, until mice were euthanized via cervical dislocation 3 hours after administration of the fourth dose. For long-term treatments, injections were administered every 48 hours with tumor growth measured 3 times weekly until sarcomas exceeded 13 mm in any dimension, at which point mice were euthanized via cervical dislocation following IACUC guidelines at Duke University. Tumor, heart, kidney, liver, brain, muscle, and plasma samples were collected and immediately snap-frozen in liquid nitrogen. For the longitudinal treatment study, sections of tumors were preserved for histological analysis.

### Histology and Microscopy

Fresh tumor samples were harvested after euthanasia, fixed in 4% PFA overnight, and preserved in 70% ethanol until paraffin embedding. Immunohistochemistry was performed in order to stain for CD31 as previously described^44^. Representative images of each H&E section were captured using a Leica DM IL LED microscope equipped with a Leica MC170HD camera with a 20× objective using LAS EZ software (Leica). Scale bars = 20 μm.

### Single-cell RNA Sequencing

Tumors were dissected and minced following the manufacturer’s protocol using MACS C tubes and the mouse Tumor Dissociation Kit (Miltenyi Biotec). After tumor dissociation, the cells were filtered through a 40 μM strainer. Red blood cells were lysed using ACK Lysing Buffer (Lonza) and washed with flow buffer made of HBSS (cat 13175-095, Gibco), 5 mM EDTA (E7899, Sigma-Aldrich), and 2.5% FBS (Gibco). Cells were washed twice more in 0.04% bovine serum albumin (BSA) in PBS, then resuspended at 1000 cells per μL. Cell suspensions were loaded on the 10x Genomics Chromium Controller Single-Cell Instrument (10x Genomics) using the Chromium Single Cell 3’ Reagent V3 Kit. Cells were mixed with reverse transcription reagents, gel beads, and oil to generate single-cell gel beads in emulsions (GEM) for reverse transcription (RT). After RT, GEMs were broken, and the single-stranded cDNA was purified with DynaBeads. cDNA was amplified by PCR and the cDNA product was purified with the SPRIselect Reagent Kit (Beckman Coulter). Sequencing libraries were constructed using the reagents provided in the Chromium Single-Cell 3’ Library Kit following the user guide. Sequencing libraries were sequenced with the Illumina Novaseq 6000 platform at the Duke GCB Sequencing and Genomic Technologies Core.

### Single-cell RNA sequencing data processing

The single cell sequencing data were processed into the gene expression tables using the pipelines from C*ell Ranger* v3.0.2 (https://support.10xgenomics.com/). Briefly, raw base call files generated by Illumina sequencers were first demultiplexed into sample-specific FASTQ files with the *cellranger mkfastq* pipeline. The FASTQ files for each sample were then aligned to the mouse reference genome (mm10) using *STAR*^75^. The aligned reads for each gene were further counted by the *cellranger count* pipeline. Quality control and filtering steps were performed to remove the low-quality cells (with fewer than 1800 genes detected) and uninformative genes (detected in fewer than 10 cells) for the downstream analyses.

### Classification of single cells into malignant and non-malignant cells

Since malignant cells typically harbor large-scale copy number alteration (*i.e.* gains or deletions of whole chromosomes or large chromosomal regions) that distinguish them from non-malignant cells ^76–79^, we performed the copy number variations (CNVs) analysis for each sample to classify the single cells into malignant and non-malignant cells. The copy number profiles were estimated based on the average expression of large sets of genes in each chromosomal region using *inferCNV* v0.99.7 (https://github.com/broadinstitute/inferCNV). A processed single-cell transcriptomic dataset from limb muscles of two healthy mouse ^42^ was served as the reference for CNVs calling. The same filtering procedures were applied to this reference dataset so that the cells with less than 1800 expressed genes, and the genes expressed in fewer than 10 cells were excluded. Genes with average expression values larger than 0.1 in reference cells were included in the following analyses. The CNVs were estimated by sorting the genes according to their chromosomal positions and using a moving average window with length 101 within each chromosome. For each sample, the cells were separated into two clusters based on the hierarchical clustering of CNV scores. We assigned the cluster with low-frequency CNVs like the reference cells as the non-malignant, while the other cluster with high-frequency CNVs was considered as the malignant.

### Classification of cell populations within non-malignant cells

The Seurat v3.0.2 (http://satijalab.org/seurat/) R package was used for the identification of non-malignant cell types. The gene expression data were firstly log-normalized and scaled with default parameters. The top 2000 most variable genes selected by Seurat were used in the principal component analysis (PCA). The first 85 principal components (PCs) selected based on the built-in jackstraw analysis were used for downstream clustering analysis and t-SNE analysis. Cell clusters were defined using FindClusters functions implemented in Seurat with default parameters and resolution=0.15. The t-SNE analysis was used to visualize the clustering results with perplexity setting to 1% of cell number whenever it was larger than 30 and learning rate setting to 1/12 of cell number whenever it was above 200, as suggested by the previous study^80^. Each cluster was annotated by comparing it’s specifically expressed genes with cell markers reported in the literature^42^ and CellMarker database^81^ (http://biocc.hrbmu.edu.cn/CellMarker/). T cells were further separated into different subtypes based on the following procedures: cells were firstly classified as CD8^+^ and CD4^+^ based on the expression levels of gene *Cd8a* and *Cd4*. T cells with *Cd8a* expression level larger than 0.1 were considered as the CD8^+^. Similarly, those with *Cd4* larger than 0.1 were considered as CD4^+^ type. While the remaining cells with both *Cd4* and *Cd8a* expression below than 0.1 were tentatively labeled as the double negative T cells. CD4^+^ T cells with the total expression level of *Foxp3* and *Il2ra* higher than 0.2 were further labeled as Tregs, while other CD4^+^ T cells were labeled as Ths. Hence the T cells were initially classified as CD8^+^, CD4^+^ Tregs, CD4^+^ Ths and double-negative T cells based on the expression level of *Cd8, Cd4, Foxp3*, and *Il2ra*. We identified the differentially expressed genes (Bonferroni-corrected p-value < 0.1) for these four groups of T cells using the FindAllMarkers function in Seurat and then performed the PCA based on these identified differential genes. The top 4 PCs selected based on jackstraw analysis were used for the next clustering analysis. We then re-clustered these T cells into 5 groups using the FindClusters function with resolution=0.1 and annotated them as the CD8^+^, Tregs, Ths, natural killer T cell (NKT) by comparing to known cell markers, as well as an unknown cell group with no discernible markers.

### Differential gene expression and pathway enrichment analysis

The Wilcoxon Rank Sum test was performed on metabolic genes to identify differences in metabolism between single cells in vehicle and digoxin treatment groups. Cohen’s d, a measure of effect size, was calculated as below to estimate the magnitude of changes in gene expression in response to Digoxin treatment.

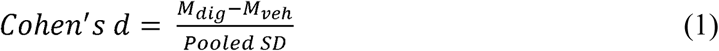

Where *M*_*dig*_ and *M*_*veh*_ is the average of each gene’s expression within a cell population from digoxin and vehicle treatment groups, respectively. The *Pooled SD*, represents the population standard deviation of a gene, is given by,

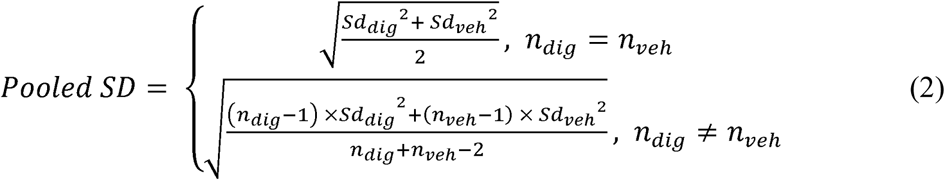

Where *n*_*dig*_ and *n*_*veh*_ are number of single cells in each population; the *Sd*_*dig*_ and *S*_*dveh*_ are standard deviations. Lists of metabolic genes and pathways were obtained from the KEGG database (https://www.kegg.jp/). The metabolic genes with *p*-value smaller than 0.01 (for malignant cells) or 0.05 (for nonmalignant cells) and the absolute value of Cohen’s d larger than 0.1 were considered statistically significant and included in the pathway enrichment analysis. The one-tailed Fisher’s exact test was used to evaluate the enrichment significance of differential metabolic genes in each metabolic pathway. GSEA analysis was performed using the javaGSEA package available at https://www.gsea-msigdb.org/gsea/downloads.jsp with default parameters. The completed differential gene sets (i.e. both metabolic and non-metabolic genes) were searched against KEGG pathways using the Metascape (http://metascape.org).

### Quantification and statistical analysis

All error bars were reported as +/− SEM with n=3 independent biological replicates and statistical tests resulting in p-value computations were obtained using a two-tailed Student’s t-test unless otherwise noted. All statistics were computed using Graphpad Prism 6 (GraphPad, http://graphpad.com/scientific-software/prism/) unless otherwise noted. * p < 0.05; **p < 0.01; ***p < 0.001.

## Data Availability

The scRNA-seq data generated during this study are available in Gene Expression Omnibus (GEO) under accession GSE149751 and can be visualized through the Single Cell Portal (https://singlecell.broadinstitute.org/single_cell/study/SCP916).

## Code Availability

Scripts reproducing the single-cell RNA-seq analysis are available at https://github.com/LocasaleLab/mouse-sarcoma-scRNA

## Author Contributions

Conceptualization, S.M.S. and J.W.L.; Computational Data Analysis, Z.X.; Animal Experiments, A.J.W., S.M.S., M.E.R., and D.G.K.; Single-cell RNA Sequencing, E.H. and S.G.G.; Metabolomics, S.M.S.; Isotope Tracing, S.M.S. and S.B.; Metabolic Flux and IC_50_ Correlations, M.V.L.; All Other Experiments, S.M.S.; Writing, S.M.S. with critical input from all authors; Supervision, S.M.S. and J.W.L.

## Acknowledgments

We thank members of the Locasale laboratory for their helpful feedback and advice. We also thank members of the Kirsch laboratory for their assistance with the mouse studies. We would also like to thank the Duke GCB Sequencing and Genomic Technologies Core for their help with RNA sequencing. Work was supported by the National Institutes of Health (F31CA224973 to SMS, R01CA193256 to JWL, F30CA221268 to AJW, R35CA197616 to DGK). Support from the Duke Compute Cluster and Data Commons Storage for computational resources are gratefully acknowledged. J.W.L serves advisory roles in Nanocare Technologies, Raphael Pharmaceuticals, and Restoration Foodworks.

## Notes

### Competing Interest Statement

Jason W. Locasale serves advisory roles in Nanocare Technologies, Raphael Pharmaceuticals, and Restoration Foodworks.

### Summary of Updates

Title, abstract, and Figure 3 revised.

