## Supplementary Information for "The Na^+^/K^+^ ATPase Regulates Glycolysis and Modifies Immune Metabolism in Tumors"

Supplementary Information for
**The Na^+^/K^+^ ATPase Regulates Glycolysis and Modifies Immune Metabolism in Tumors**

Sanderson SM, Xiao Z et al.

Supplementary Figure 1.
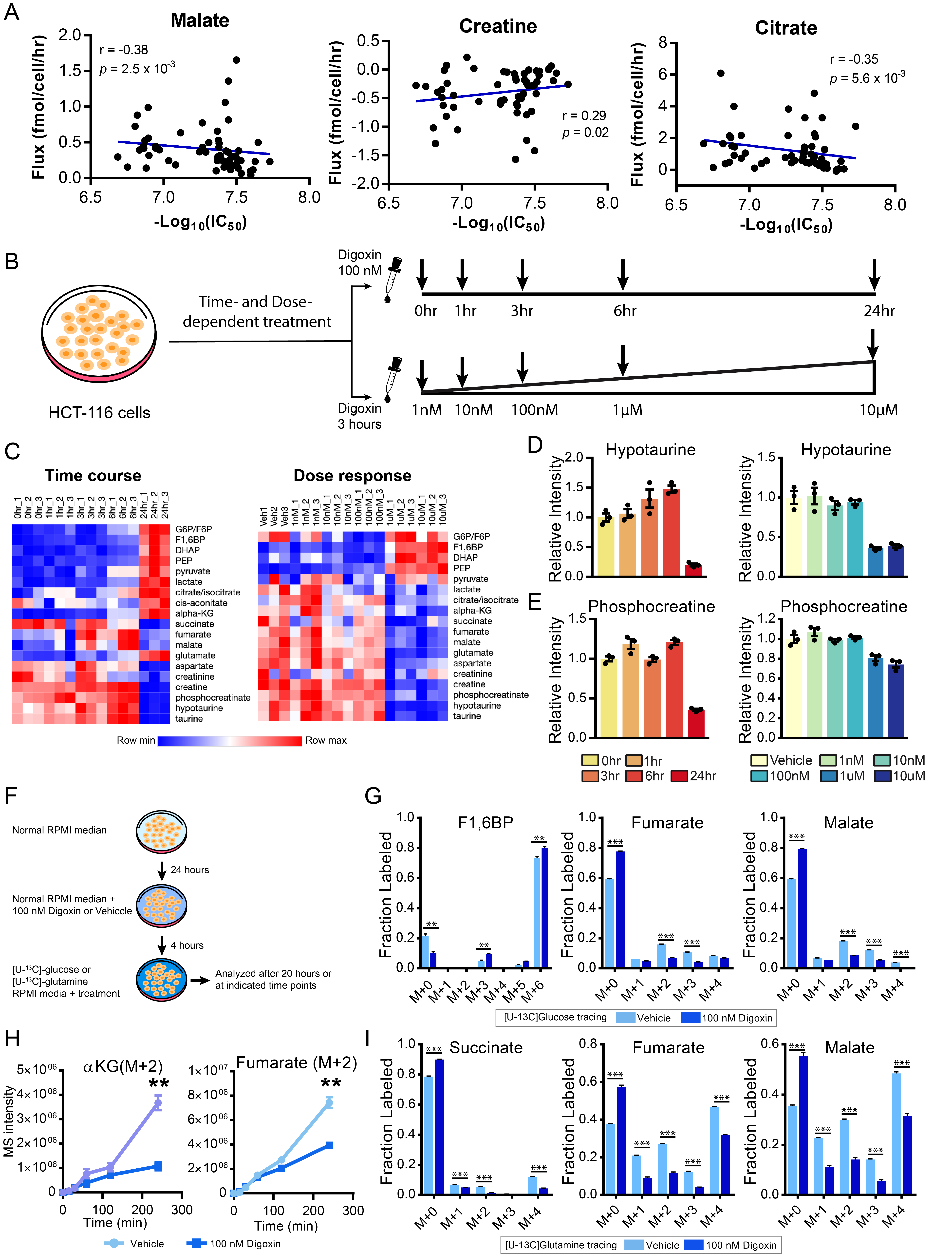

#### Supplementary Figure 1. Digoxin induces broad alterations in central carbon metabolism.

(A) Scatter plot showing correlation of basal metabolic flux with digoxin IC_50_ across NCI-60 cell line panel. Each dot represents a cell line from the NCI-60 cell panel.

(B) Diagram of time course and dose-response experiment design.

(C) Heatmaps of time- and dose-dependent alterations of metabolites in central carbon metabolism.

(D) Relative intensity of hypotaurine at each time point and dose level.

(E) Same as in (D) but for phosphocreatine.

(F) Diagram showing the design of both steady-state and kinetic labeling in [U-^13^C] glucose and [U-^13^C] glutamine tracing experiments with HCT-116 cells.

(G) Fractional abundance of each [U-^13^C] glucose labeled isotopologue to the sum of all isotopologues of the indicated metabolites. F1,6BP: fructose 1,6 bisphosphate.

(H) Relative intensities of the [U-^13^C] glucose isotopologue (M+2) of α-ketoglutarate (αKG) and fumarate at indicated time points.

(I) Fractional abundances of [U-^13^C] glutamine labeled isotopologues of TCA cycle intermediates as in (G).

Data in (D), (E), (G), (H) and (I) are expressed as mean ± SEM of n = 3 biological replicates. * *p* < 0.05; ***p* < 0.01; ****p* < 0.001 as determined by Student’s t-test.

Supplementary Figure 2.
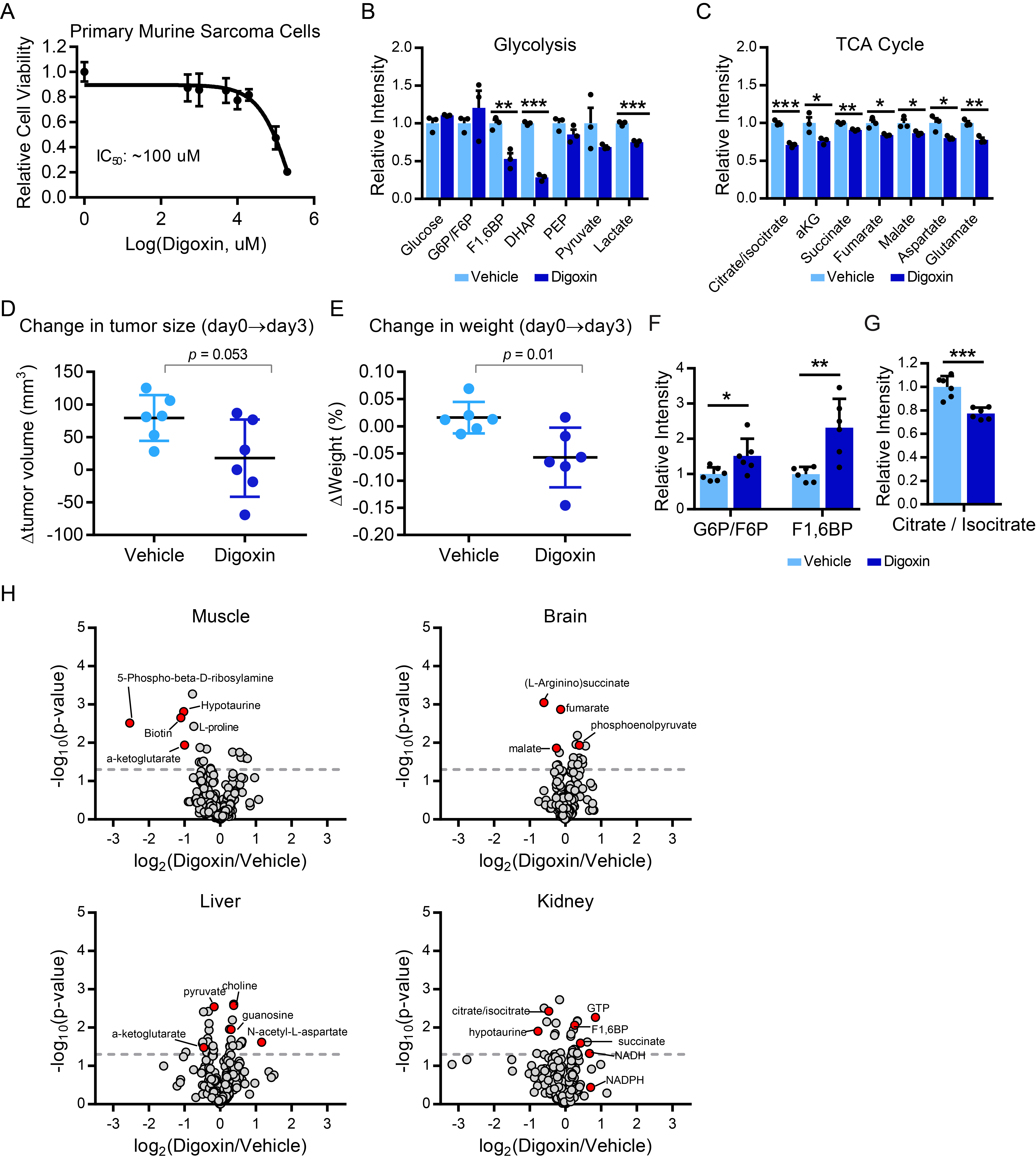

Supplementary Figure 2.
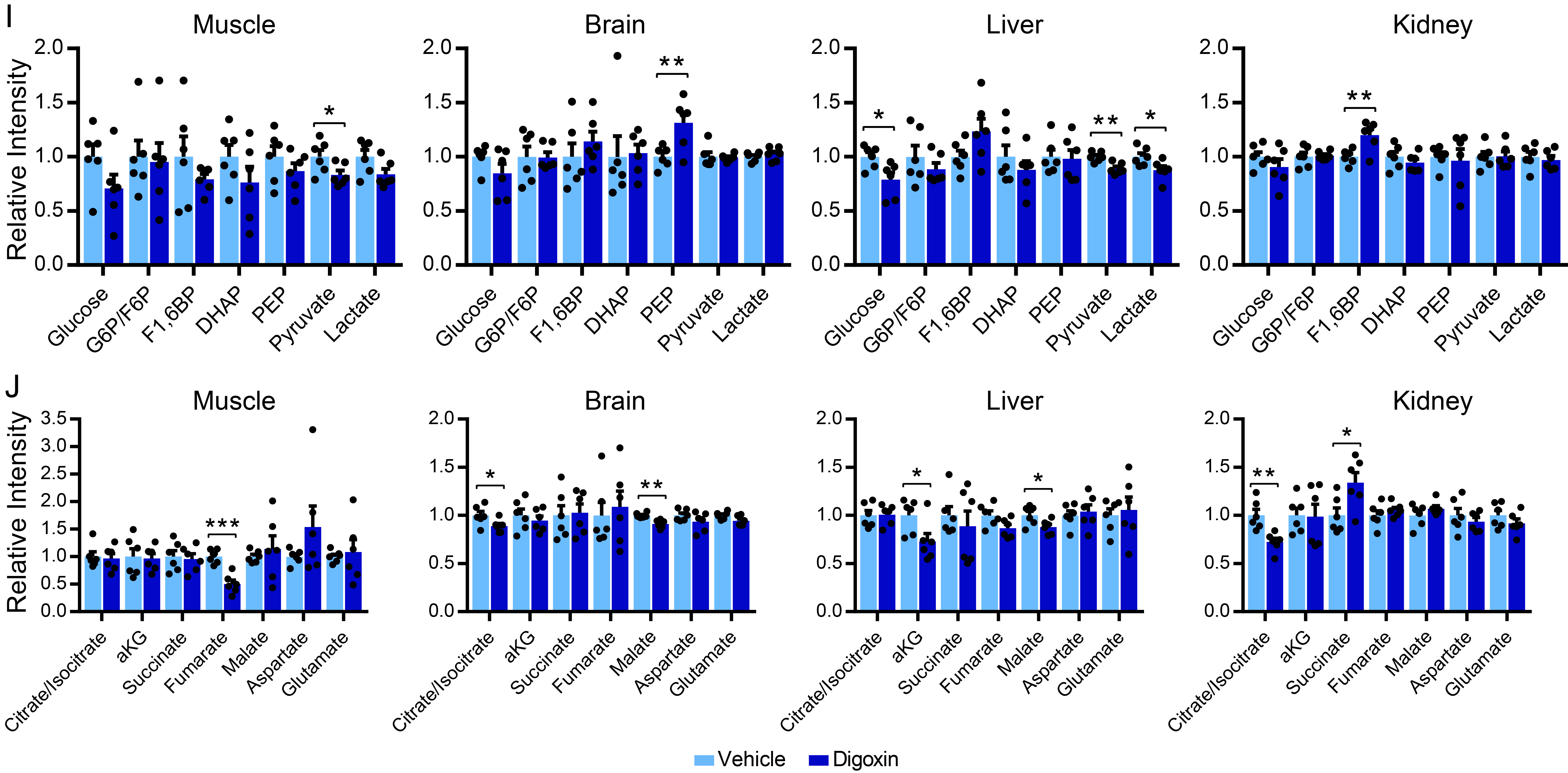
Digoxin impacts metabolic processes in murine cells of diverse origin.

(A) Determination of digoxin IC_50_ in cultured primary murine sarcoma cells.

(B and C) Relative intensities of glycolytic (B) and TCA cycle (C) intermediates in cultured sarcoma cells after 48 hours of vehicle or 100 μM digoxin treatment.

(D) The difference in tumor volume between day 0 and day 3 in vehicle and digoxin treatment groups.

(E) The difference in body weight between day 0 and day 3 in vehicle and digoxin treatment groups. Other than alterations to body weight, no other behavioral toxicities were observed.

(F) Relative intensities of upper glycolytic intermediates in cardiac tissue with vehicle and digoxin treatment. N=6.

(G) Same as in (F) but for TCA intermediate citrate/isocitrate. N=6.

(H) Volcano plot of *P*-value (log10 scale) vs. fold change (log2 scale) for metabolite levels of muscle, brain, liver, and kidney tissues between vehicle- and digoxin-treatment.

(I) Same as in (B) but for glycolytic intermediates from muscle, brain, liver, and kidney tissues.

(J) Same as in (C) but for TCA cycle intermediates from muscle, brain, liver, and kidney tissues.

Data in (A)­—(C) are expressed as mean ± SEM of n=3 biological replicates; Data in (D) and (E) are expressed as mean ± SD of n=6 biological replicates; Data in (I) and (J) are expressed as mean ± SEM of n=6 biological replicates. * p < 0.05; **p < 0.01; ***p < 0.001 as determined by Student’s t-test.

### Supplementary Figure 3.

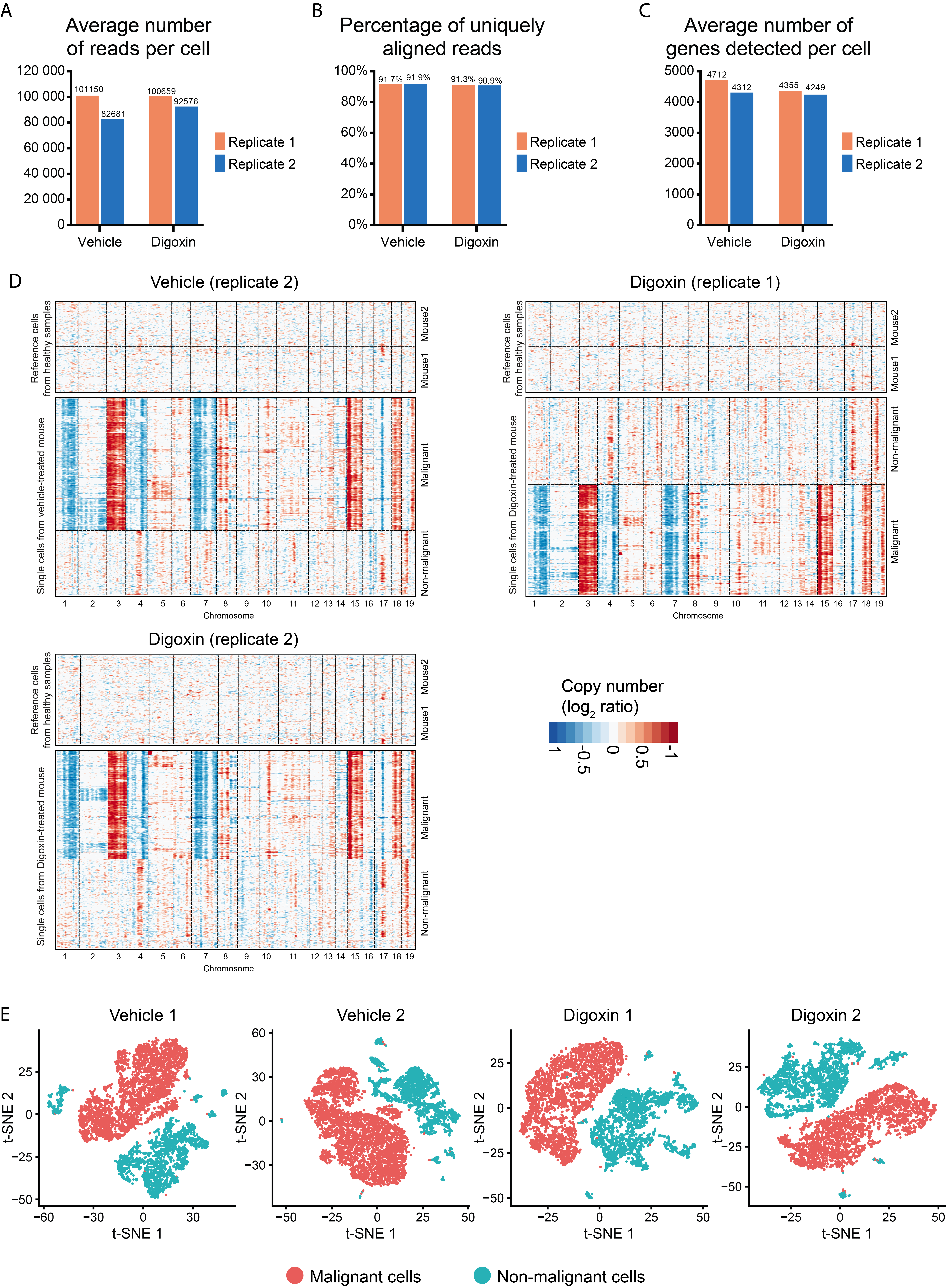

#### Supplementary Figure 3. Summary of Single-cell RNA sequencing data and copy number variation analysis.

**(A)** Average number of sequencing reads captured per cell for samples treated with vehicle and digoxin.

**(B)** Average percentage of uniquely aligned reads per cell for samples treated with vehicle and digoxin.

**(C)** Average number of detected genes per cell for samples treated with vehicle and digoxin.

**(D)** Chromosomal landscape of large-scale copy number variations (CNVs) of single cells from normal mouse muscle tissue (top) and sarcoma tumors treated with vehicle and digoxin. Y-axis represents individual cells. Amplifications (red) and deletions (blues) were inferred by averaging gene expression over 100 genes stretch on each chromosome (columns).

**(E)** t-SNE plots of scRNA-seq data demonstrating distinct clustering between malignant and non-malignant cells based on CNV results from (D).

### Supplementary Figure 4.

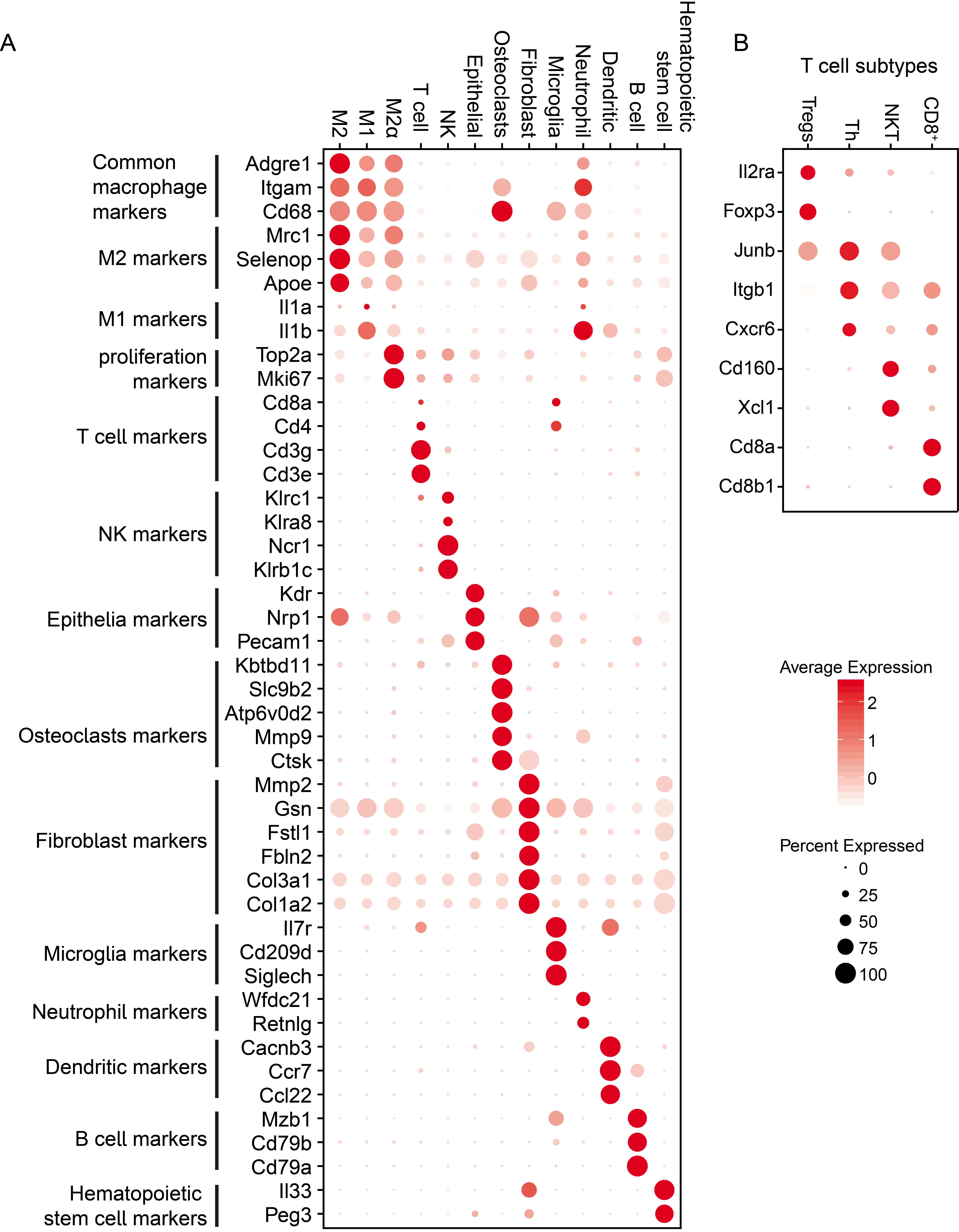

#### Supplementary Figure 4. Cell-type assignment of single-cell populations using the expression of marker genes

(A) Bubble plot comparing the scaled expression of marker genes across non-malignant cell clusters. Bubble size refers to the proportion of cells in a cluster expressing a gene, and the color represents the average scaled expression within a cluster. The source of marker genes for assigning the cell types are described in ‘Methods’. M1: type I macrophages; M2: type II macrophages; M2α: highly proliferation M2; NK: Natural killer cells.

(B) Similar as in (A) but for T cell subpopulations. Tregs: Regulatory T cells; Th: T-helper cells; NKT: Natural killer T cells.

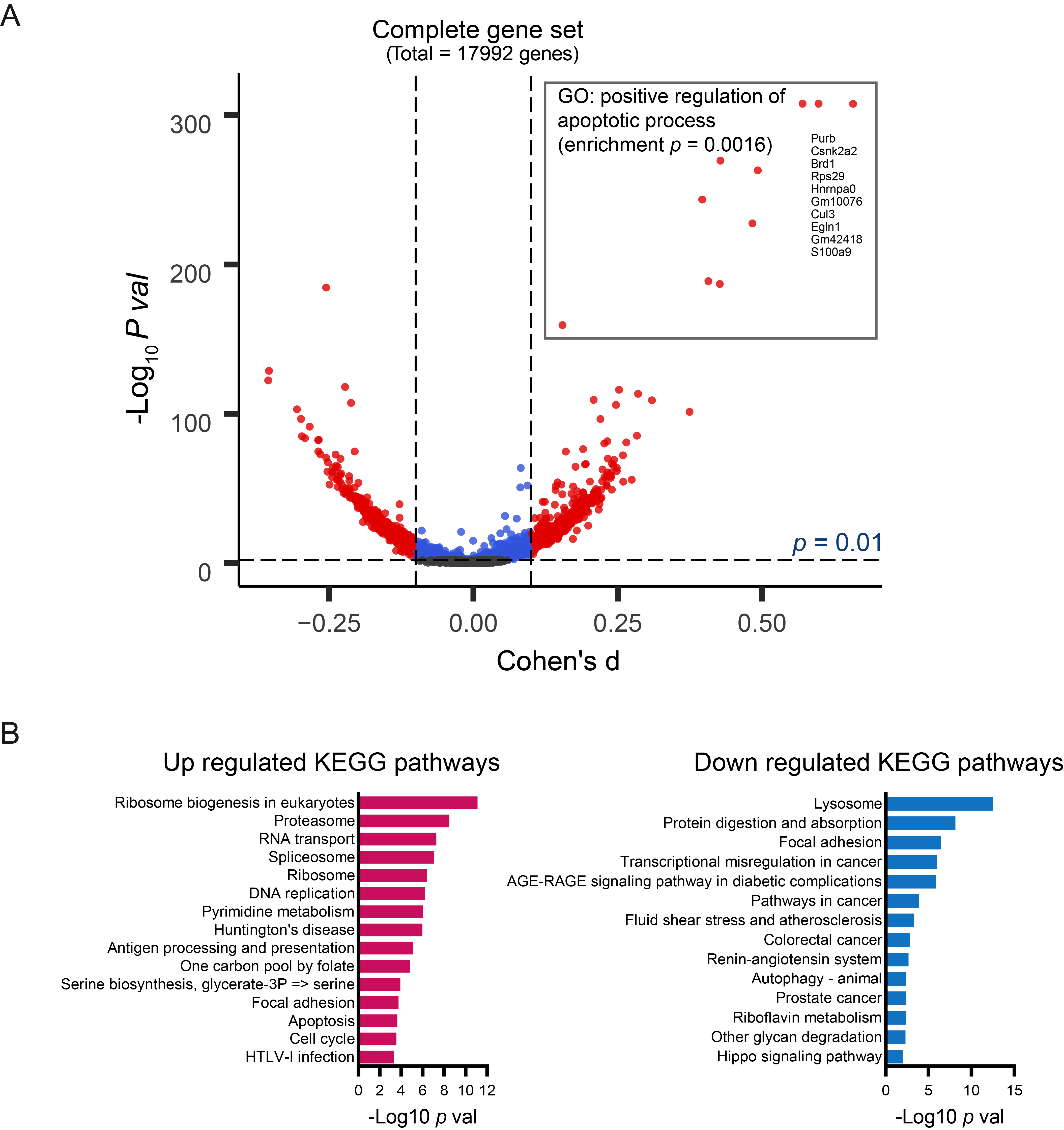

#### Supplementary Figure 5. Regulation of gene expression in malignant cells suggests the association between digoxin treatment and cell apoptosis.

(A) Volcano plot showing the gene expression difference (measured by Cohen’s d) for all genes between vehicle and digoxin treatment versus the -log_10_ *p* value for that difference. The differential genes above a significance threshold of p value < 0.01 and the absolute value of

Cohen’s d > 0.1 are labeled in red. The gene symbols of top 10 up-regulated genes in digoxin treatment group and their associaated GO term are annotated in the gray rectangular box.

(B) The enriched pathways determined by KEGG enrichment analysis using significantly upregulated and downregulated genes indicated in (A).

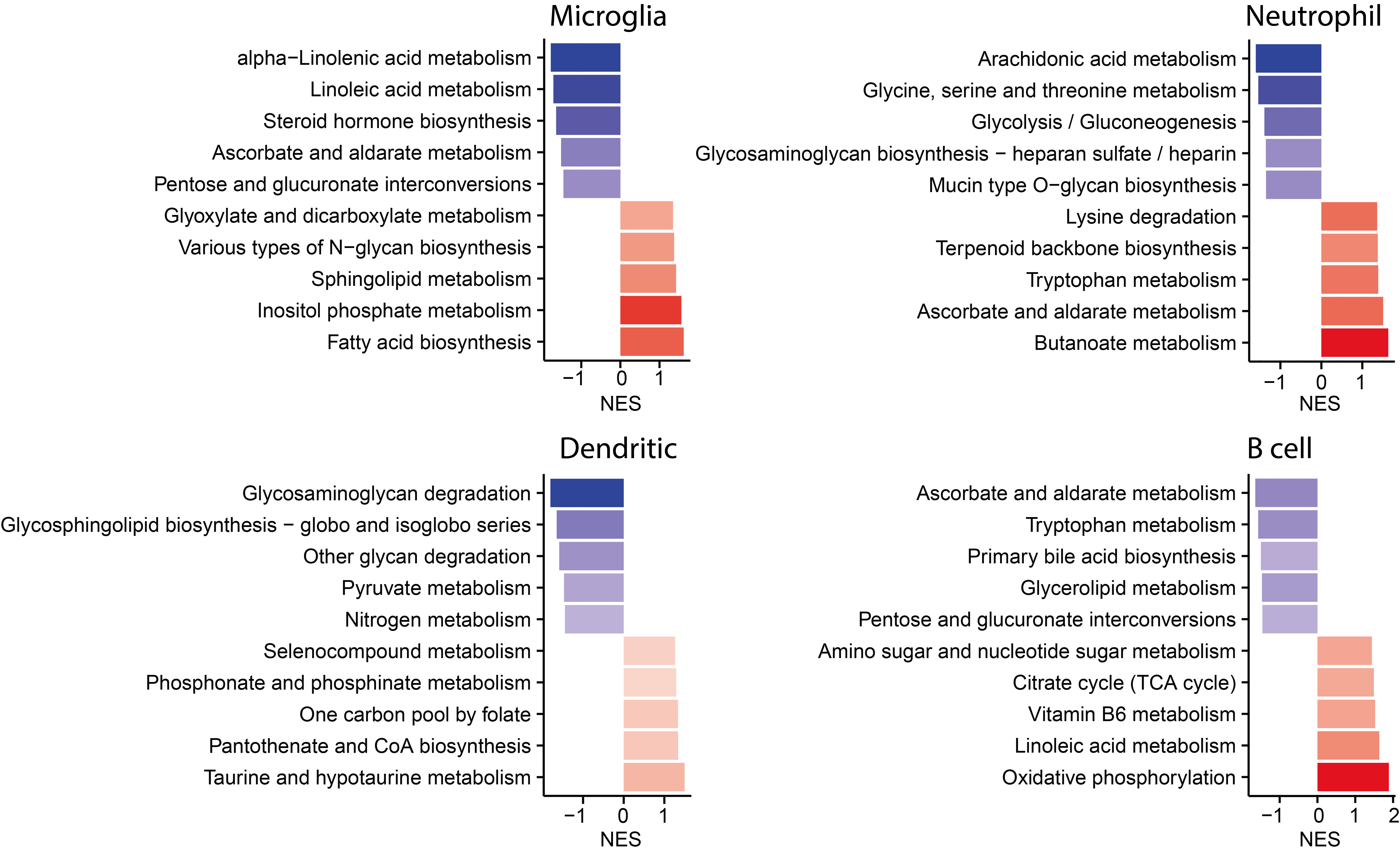

#### Supplementary Figure 6. Metabolic pathways enriched in digoxin- versus vehicle-treated sarcoma tumors.

Top 10 pathways enriched in GSEA analysis using metabolic genes. Red: higher in digoxin treatment; Blue: higher in vehicle treatment. NES: normalized enrichment score. Each bar is colored by p-value, with warmer colors indicating much higher significance.

### Supplementary Figure 7.

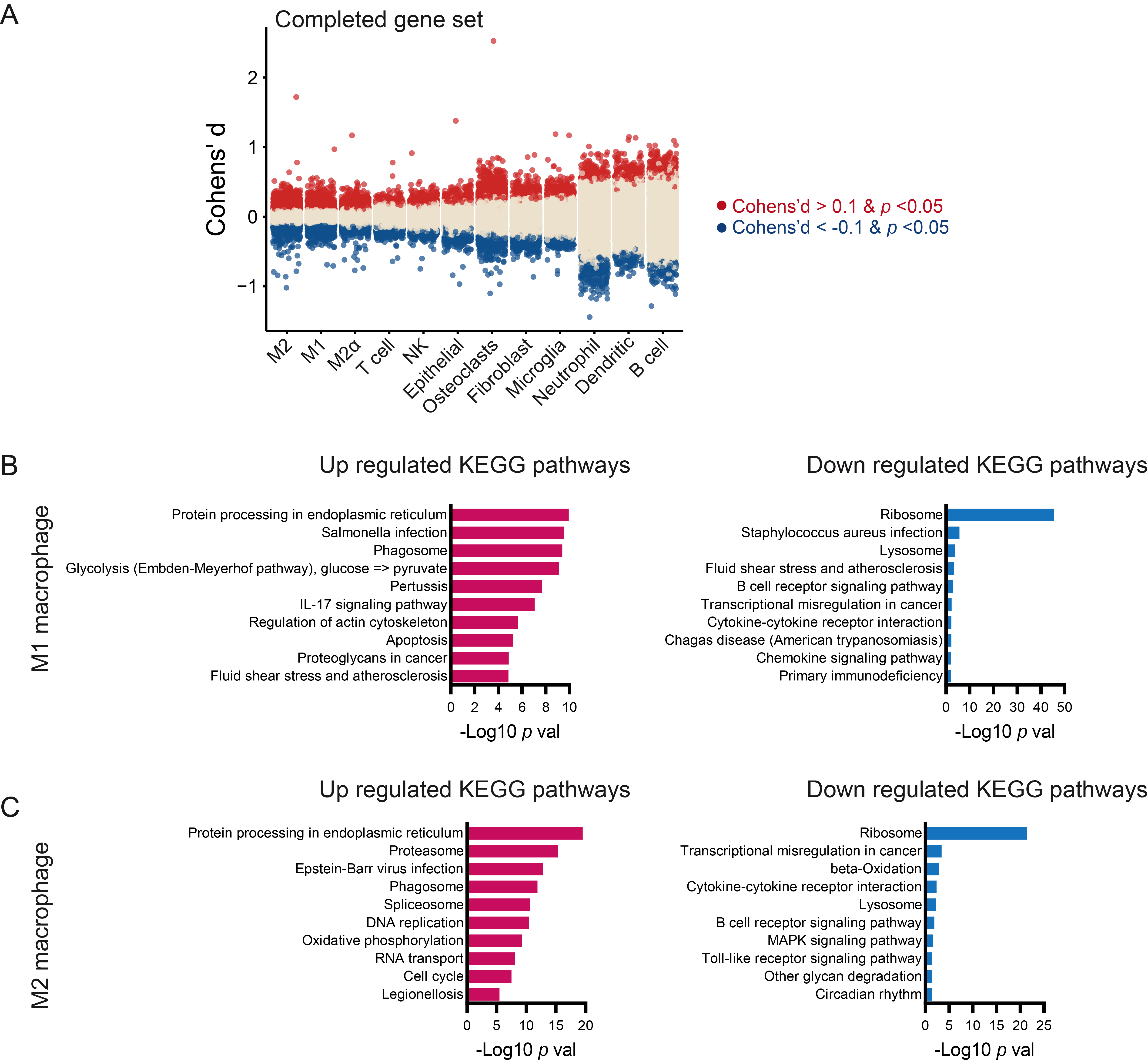

#### Supplementary Figure 7. Regulation of gene expression in macrophage cells suggests the association between digoxin treatment and cell activation.

(A) Distribution of Cohen’s d for gene transcripts from full transcriptome in each cell population. The upregulated and downregulated genes in digoxin treatment are marked in red and blue respectively.

(B) Top 10 enriched pathways determined by KEGG enrichment analysis using significantly upregulated and downregulated genes in M1 macrophages.

(C) Same as in (B) but for M2 macrophages.
